# Linguistic coupling between neural systems for speech production and comprehension during real-time dyadic conversations

**DOI:** 10.1101/2025.02.14.638276

**Authors:** Zaid Zada, Samuel A. Nastase, Sebastian Speer, Laetitia Mwilambwe-Tshilobo, Lily Tsoi, Shannon Burns, Emily Falk, Uri Hasson, Diana Tamir

## Abstract

The core use of human language is communicating complex ideas from one mind to another in everyday conversations. In conversations, comprehension and production processes are intertwined, as speakers soon become listeners, and listeners become speakers. Nonetheless, the neural systems underlying these faculties are typically studied in isolation, using paradigms that cannot fully engage our capacity for interactive communication, and with indirect measures of similarity. Here, we used an fMRI hyperscanning paradigm to measure neural activity simultaneously in pairs of subjects engaged in real-time, interactive conversations. We used contextual word embeddings from a large language model to quantify the linguistic coupling between production and comprehension systems within and across individual brains. We found a highly overlapping network of regions involved in both production and comprehension spanning much of the cortical language network. Our findings reveal that shared representations for both processes extend beyond the language network into areas associated with social cognition. Together, these results suggest that the specialized neural systems for speech perception and production align on a common set of linguistic features encoded in a broad cortical network for language and communication.

## Introduction

Everyday language relies on two fundamental processes: production and comprehension (e.g., speaking and listening). While these may seem different and are often studied separately, there is evidence for shared representations and mechanisms between them (Gambi & Pickering, 2017; Giglio et al., 2022; Hu et al., 2023; Menenti et al., 2012; Papathanassiou et al., 2000; Silbert et al., 2014). However, to truly test the overlap between these neural systems, we need to explicitly model their linguistic representations and study dyads in real-time interactive conversation who engage in both speaking and listening. Recent advances allowed us to address these points by using large language models (LLMs) as explicit computational models of brain activity during real-time conversations using fMRI hyperscanning.

Prior studies on the similarity between production and comprehension neural processes have only indirectly tested their overlap. These studies typically use different tasks for speaking and listening and then compare whether the same brain region is activated for both. For example, Awad and colleagues (2007) localized brain regions for comprehension using a contrast between listening to regular speech versus rotated speech; and regions for production by contrasting activity during counting versus speaking. Contrast-based approaches may find the same region involved in both processes (Hu et al., 2023), but this does not inform us of the similarity between their representations. Some studies used a two-brain approach, such as speaker-listener coupling as a data-driven test of similarity (Silbert et al., 2014; Stephens et al., 2010). While others rely on an adaptation paradigm to probe for syntactic similarity (Menenti et al., 2011; Segaert et al., 2012). However, these methods are fundamentally content-agnostic—they cannot tell us “what” is shared between brains or processes. To quantify linguistic representations shared between communicators, we need an explicit model of linguistic features (Zada et al., 2024). Using explicit features would allow us to quantitatively compare the underlying representations across production and comprehension.

Encoding models have recently been used to quantify linguistic features encoded in neural activity during passive language comprehension (de Heer et al., 2017; Huth et al., 2016; Wehbe et al., 2014), and less commonly, to language production (Cai et al., 2025; Goldstein et al., 2025; Yamashita et al., 2025). LLMs have been shown to have similarity judgments comparable to humans (De Deyne et al., 2019; Levy et al., 2015), and have internal linguistic representations that are more similar to the human brain than any other model, during language comprehension (Caucheteux et al., 2023; Goldstein et al., 2022; Heilbron et al., 2022; Schrimpf et al., 2021). By leveraging the rich linguistic representations from LLMs, encoding models can identify the components of neural activity that encode linguistic features. These methods have begun to lend further support for the integrated view of neural representations for production and comprehension both within subjects (Cai et al., 2025; Goldstein et al., 2025; Yamashita et al., 2025), and across subjects (Zada et al., 2024) during real-time dialogue. For example, Zada and colleagues (2024) used contextual embeddings from an LLM as an explicit, shared model mediating speaker-listener neural coupling during real-time conversations in electrocorticography. Most studies using LLM embeddings for encoding analyses have only been applied to comprehension, and those that include production only do so while recording from one subject.

In real-time dyadic conversations, production and comprehension are contemporaneous and interleaved. Production is often spontaneous and comprehension must be proactive: as listeners must be ready to respond in a relevant way as they process the incoming speech (Grice, 1975; Redcay & Schilbach, 2019). Conversation is unique in that it requires coordination in the form of interactive alignment (Pickering & Garrod, 2004) and agreement on meaning through common ground (Brennan & Clark, 1996; Clark & Brennan, 1991; Wilkes-Gibbs & Clark, 1992). Conversation is also arguably the most fundamental setting of language use. It is universal to human societies, does not require specialized skills (e.g., literacy) or technologies (e.g., telephones) (Clark, 1996), and allows people to go well beyond simple stimulus-response signaling to share and shape each others’ representational thought through language. However, previous research has decoupled production and comprehension, using separate tasks and stimuli for each process, controlled paradigms (e.g., rehearsed speech, covert production), and isolated linguistic contexts. This raises the question of whether these paradigms fully capture the neural systems for real-time, interactive conversation (Hasson & Honey, 2012; Nastase et al., 2020).

Conversational language is fundamentally a social process. Previous work has investigated production–comprehension coupling across subjects using a sequential, asynchronous protocol: first recording a subject speaking and then playing the speech back to multiple listeners at a later time (Chang et al., 2023; Jiang et al., 2012; Kinreich et al., 2017; Liu et al., 2022; Nguyen et al., 2022; Silbert et al., 2014; Stephens et al., 2010; Zadbood et al., 2017). Using content-agnostic measures, these studies find that during communication, the speaker’s neural activity is coupled to that of the listeners in regions associated with both comprehension and production. Moreover, the strength of speaker-listener coupling is related to outcome measures of comprehension. Hyperscanning paradigms, where researchers simultaneously measure neural processes during dyadic social interactions using two MRI scanners, are uniquely suited to studying language usage in interactive social settings (Babiloni & Astolfi, 2014; Czeszumski et al., 2020; Montague et al., 2002; Nam et al., 2020; Redcay & Schilbach, 2019; Speer et al., 2024; Tsoi et al., 2022; Wheatley et al., 2019), and can inform the extent of shared neural mechanisms and speaker-listener alignment (Hasson et al., 2012; Schoot et al., 2016). Paradigms of this kind can extend asynchronous work on the neural systems for production and comprehension as they couple in real-time conversations across brains.

In this paper, we developed an fMRI hyperscanning paradigm to simultaneously measure whole brain activity in dyadic pairs of subjects engaged in free-form, interactive conversations across a range of prompted topics. We used encoding models to map the cortical areas representing linguistic content during spontaneous speech production and comprehension. Within subjects, we found overlapping cortical systems for production and comprehension: both systems depend strongly on common representations and only partially on specialized representations. Then, using the real-time, dyadic conversations paradigm, we compute model-based coupling across the speaker’s and listener’s brains via the LLM embeddings. We found that model-based speaker-listener coupling engages areas associated with social cognition. Through model-based LLM encoding analysis of whole brain fMRI neural signals, we gain valuable insights into how successful conversations depend on shared language representations between production and comprehension across various cortical regions.

## Results

We aimed to model production and comprehension processing within and between brains during free-form, turn-based conversations. We used hyperscanning to collect simultaneous fMRI data in 30 dyads (60 subjects) as they freely discussed ten topics across five ∼6 min runs (Figure 1A) (Speer et al., 2024). Topic prompts were presented as a starting point, but each dyad was free to pursue the discussion differently, resulting in 30 unique conversations (Table S1). We hypothesized that we could use a computational model to quantitatively evaluate the overlap between production and comprehension (Figure 1B). To characterize the linguistic content in the BOLD signal, we explicitly represented the language stimuli with several different feature spaces: confound variables (e.g., word rate), spectral acoustic features, phonemic articulatory features, and word embeddings extracted from GPT-2 (Radford et al., 2019). Then, we used banded ridge regression to estimate a linear mapping from the model features onto the BOLD activity at each vertex (Dupré La Tour et al., 2022; Huth et al., 2016; Naselaris et al., 2011; Nunez-Elizalde et al., 2019) (Figure 1C). To evaluate the models, we correlated the model-predicted and actual BOLD time series for left-out runs for each feature space and for production or comprehension time points separately (Figure 1D). Finally, we averaged the model performance correlations across subjects for all analyses. Statistically, we evaluated the average using a one-sample t-test, correcting for multiple comparisons over all ∼75k cortical vertices. To summarize our results, we averaged the encoding performance across vertices within 11 regions of interest (ROIs) spanning an extended language network, from low-level auditory and articulatory areas to high-level semantic areas: early auditory cortex (EAC), posterior and anterior superior temporal gyrus (pSTG, aSTG), inferior and middle frontal gyri (IFG, MFG), somatomotor cortex (SM), supplementary motor area (SMA), frontal opercular (FOP), intraparietal sulcus (IPS), temporoparietal junction (TPJ), and posterior medial cortex (PMC).

**Figure 1.**
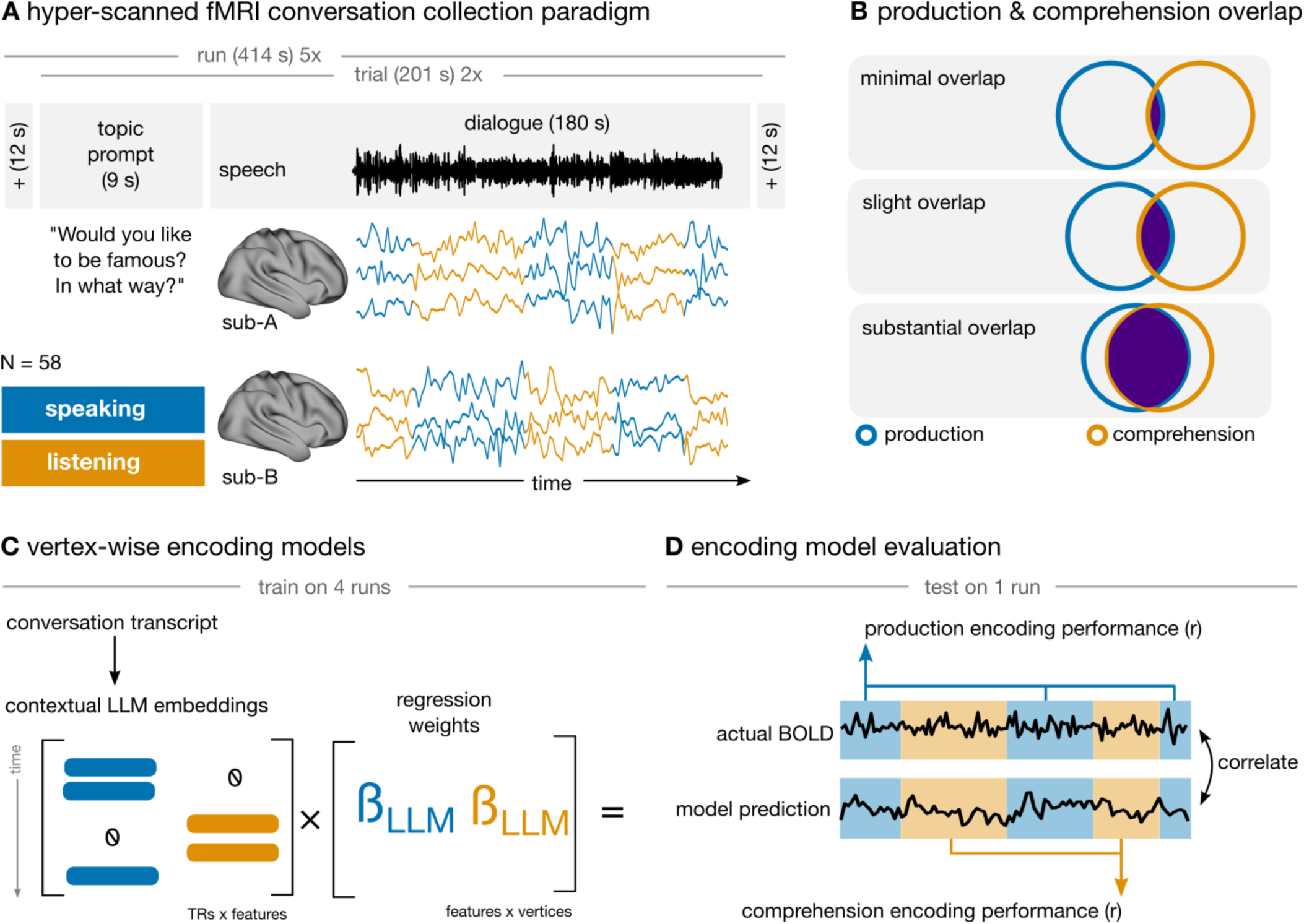
Data collection and modeling framework. (**A**) We collected fMRI data simultaneously from pairs of subjects as they engaged in interactive, prompted conversations. (**B**) We aimed to quantify the overlap of production and comprehension processes using an explicit computational model. (**C**) We extracted LLM word embeddings from each conversation transcript and used them as encoding model features to predict the subject’s brain activity. We split the regressors into separate time series for production (blue) and comprehension (orange). (**D**) Finally, we evaluated the performance of production and comprehension time points separately in a held-out test run of different conversations.

### Contextual embeddings capture both production and comprehension

We first validated that we can successfully model brain activity during spontaneous production and comprehension in our hyperscanning paradigm. To do so, we built two models to quantify linguistic processing and to measure the cortical overlap between production and comprehension. In one, we constrained the model to learn one set of shared weights for production and comprehension for all feature spaces. In this model, a vertex must code for both processes with the same functional tuning (i.e., shared weights) to be well predicted. In the second model, we split all regressors into separate sets for production and comprehension, allowing the model to learn separate weights for each process (Figure 1C). We treat the confound, acoustic, and phonemic feature sets as nuisance variables and report only the LLM contextual embedding performance. We first inspect the performance of the second, more flexible model, which we expect to outperform the unified constrained model.

Using the more flexible model with separate weights for production and comprehension, we found significant within-subject encoding performance throughout the core language network: STG, IFG, and MFG for speech production and speech comprehension (Figure 2A). Moreover, encoding performance extended bilaterally to higher-level regions like TPJ and PMC. We found considerable spatial overlap between encoding performance for production and comprehension—i.e., vertices well predicted during production are also likely to be well predicted during comprehension (*r* = 0.712, *p* < 1e-5). To quantify whether production and comprehension encoding rely on shared or divergent weights, we compared the performance of the shared-weights and separate-weights models. We found that across all 11 ROIs, a large proportion of encoding performance can be attributed to shared functional tuning rather than idiosyncratic production- or comprehension-specific variance (Figure 2B). Peripheral regions for speech perception (EAC) and speech articulation (SM) showed the largest divergence, but the shared-weights model still recovered over half the performance of the separate-weights model. These results suggest that cortical activity during both production and comprehension keys to similar features captured by the LLM embeddings.

**Figure 2.**
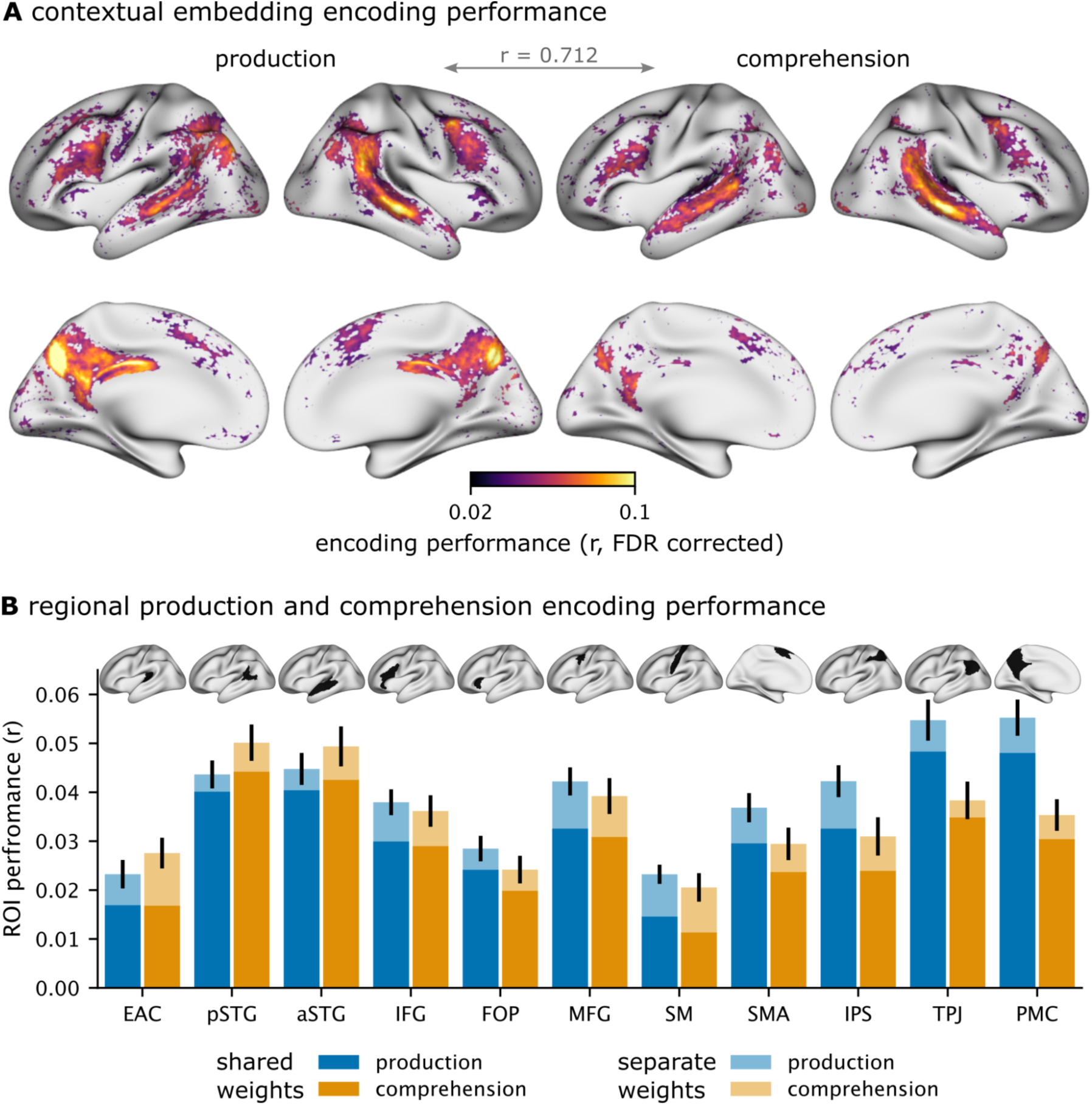
Within-subject speaking and listening encoding performance. (**A**) Encoding model performance of the separate-weights model for production and comprehension relative to the control feature spaces. (**B**) We summarized the un-thresholded encoding performance of the shared- and separate-weights models in 11 ROIs spanning the extended language network, averaged across left and right hemispheres (see Methods).

We observed several qualitative differences across tasks, regions, and hemispheres. First, overall encoding performance appears higher in right STG and TPJ than in the left-hemisphere homologs. Second, overall encoding performance appears stronger for production, especially in bilateral PMC and right TPJ. Third, encoding performance for comprehension appears stronger and more bilateral in STG than in production. Despite these differences, the overall encoding performance suggests that LLM embeddings provide a rich basis for modeling linguistic encoding throughout much of the cortex.

### Story-listening comprehension shares a subset of linguistic features with interactive production and comprehension neural systems

In addition to the hyperscanning conversations paradigm, we recorded participants as they listened to a 13-minute story in a separate scanning session. This presented a unique opportunity to compare linguistic processing during spontaneous production, (inter)active comprehension, and non-interactive comprehension. Specifically, we aimed to test the shared processing between passive listening and active comprehension and production. To do so, for each subject, we estimated a comprehension encoding model using the story data only and then evaluated the fitted model on the subject’s conversation data. We extracted the same four feature spaces from the story, and evaluated the model performance similarly to the conversation models.

We found significant within-subject generalization performance from the passive listening paradigm to the conversational paradigm for production and comprehension (Figure 3). Generalization to conversational comprehension was stronger than to production. However, both were lower than when training on conversational data, capturing only a portion of the variance as training on conversations— even when equating their training data (Figure S1). Training on conversation data resulted in an increase of +41% in average encoding performance for comprehension and an increase of +49% for production. A paired t-test found a significant difference (*p* < 0.00212) between subjects’ average vertex encoding performance when training on conversation or story. Generalization performance was more bilateral than performance based on the conversational paradigm. An overlapping set of regions was well predicted, particularly STG during comprehension and PMC during production. Notably, generalization was poorer for IFG and MFG compared to temporal regions. Though incomplete, generalization from passive comprehension to both production and comprehension in a conversation context provides further evidence for a common subset of linguistic features that span both processes, while still highlighting the boost in these systems during active, naturalistic communication.

**Figure 3.**
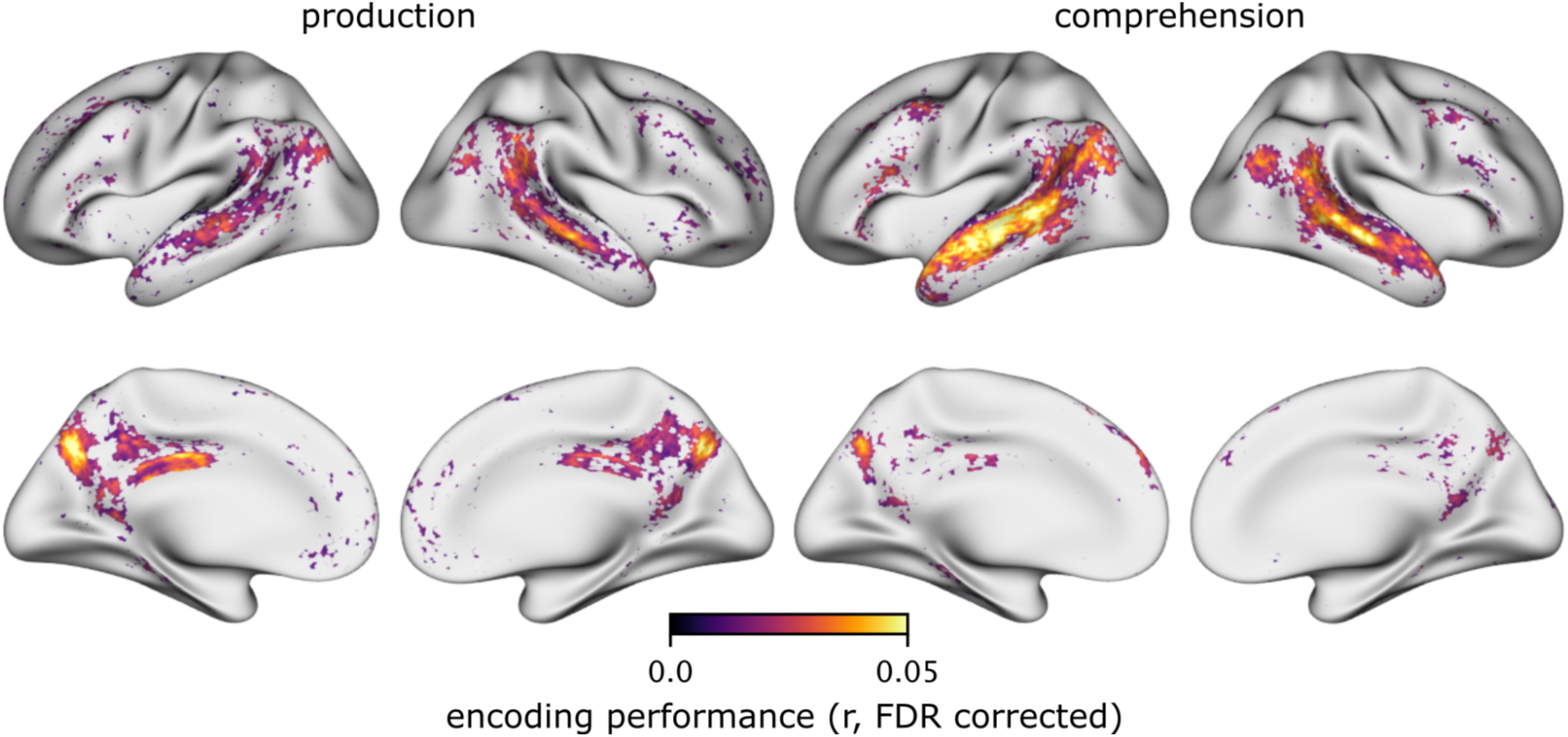
Encoding models trained on passive listening partially generalize to neural responses during conversations. Participants passively listened to a 13-minute story in a separate scanning session before the hyperscanning procedure. We estimated encoding models using the same four feature spaces from this passive listening-only dataset. Then we evaluated how well they generalize to data acquired during conversational production and comprehension conversations. Here we present only the performance of the contextual embedding feature space, after testing for significance and correcting for multiple comparisons. We found significant encoding performance in STG and PMC, which is significantly weaker than when training on conversations.

### Contextual embeddings outperform other features of speech and language

Our modeling framework allows us to test different hypotheses about features of brain activity during production and comprehension by comparing the performance of different models. So far, we have only reported the performance of the contextual word embeddings from a pre-trained language model. However, we can decompose the joint model performance into the relative contribution from each feature space. Here, we report the performance of each feature space and then use a variance partitioning analysis to compute the unique variance predicted by the contextual LLM embeddings.

We found that during both production and comprehension, the contextual LLM embeddings outperformed all other feature spaces regarding correlation strength and cortical coverage (Figure 4A, Figure S2). Among the lower-level control feature spaces, we observed that the acoustic features were the most predictive, especially in EAC and STG. In contrast, the articulation band was least predictive throughout all regions (likely due to collinearity with the better-fitting acoustic space). Moreover, the confound variables were most predictive in SM, EAC, and aSTG. These regions are likely to exhibit large signal fluctuations between speech production or comprehension and are more susceptible to regressors such as word rate.

**Figure 4.**
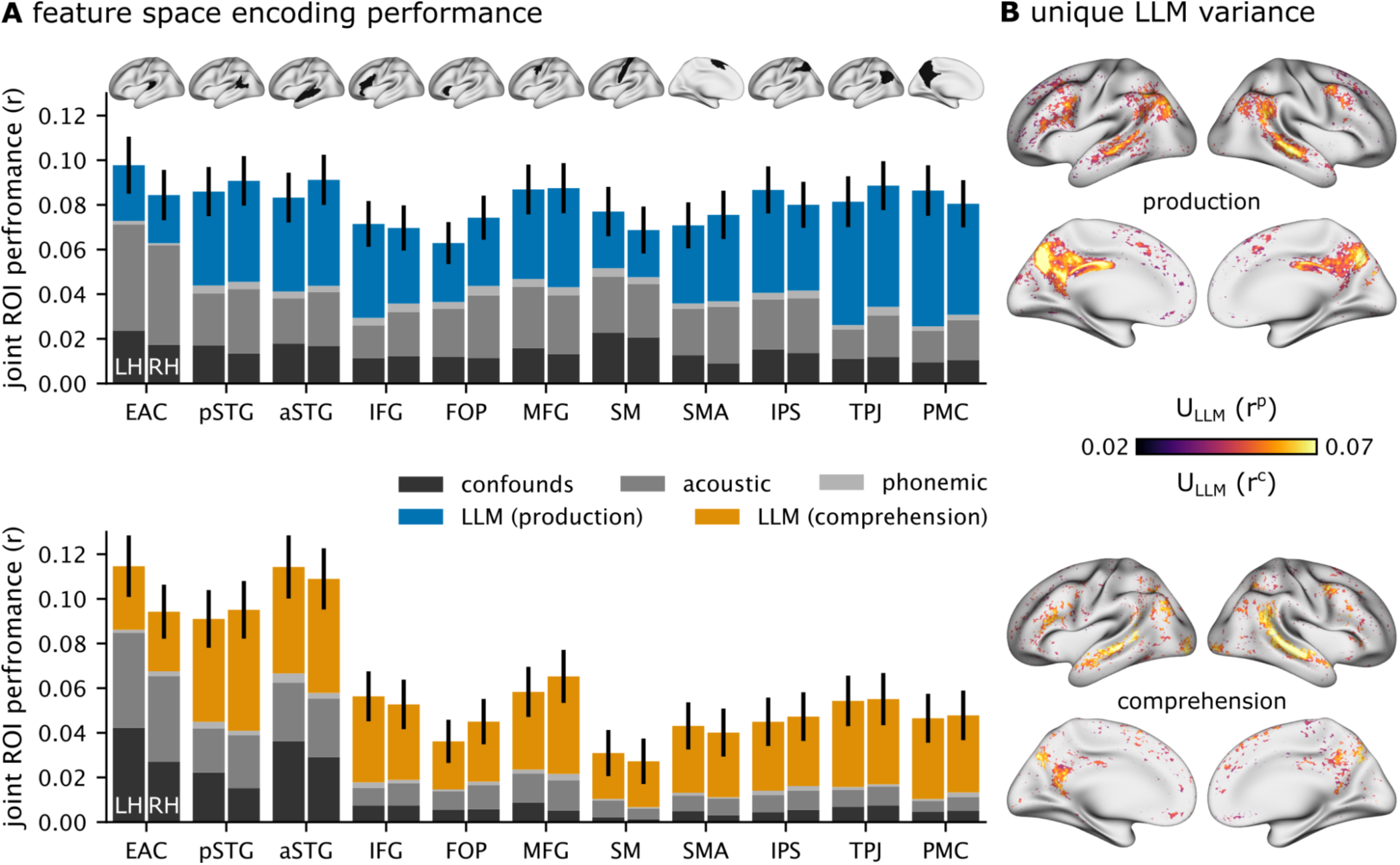
Model comparison and variance partitioning. We compared the variance explained by LLM embeddings with other linguistic feature spaces. (**A**) The joint encoding performance of the full model was decomposed into the contribution of each space separately for production and comprehension. (**B**) We performed a variance partitioning analysis within subjects to quantify the unique contribution of LLM word embeddings. We trained one full encoding model with all features and a nested model with all features, excluding the LLM word embeddings. Then, we subtracted the nested model performance from the full model to quantify the unique variance explained by the LLM embeddings.

Next, we performed a variance partitioning analysis (de Heer et al., 2017; Lee Masson & Isik, 2021; Lescroart et al., 2015) to isolate the unique variance explained by the contextual LLM embeddings. We use hierarchical regression to compare a full model with all features and a nested model excluding the features of interest. In this analysis, the full model is composed of the LLM contextual embeddings (L), acoustic (A), and articulatory phonemic (P) features, resulting in encoding performance R_L,A,P_. The nested model is the same, except that it excludes the LLM contextual embeddings from the predictors. Therefore, the unique contextual variance can be calculated as U_L_ = R_L,A,P_ – R_A,P_. The contextual embeddings accounted for unique variance bilaterally across all previously reported brain regions (Figure 4B). Together, these results suggest that while part of the variability in brain activity can be predicted by acoustic speech features, the contextual word embeddings of LLMs provide unique predictive power, especially in higher-order regions.

### Model-based brain-to-brain coupling between conversational partners

When two people converse, we expect their brain activity to align along certain shared features between speech production and comprehension (Hasson et al., 2012; Silbert et al., 2014; Stephens et al., 2010; Zada et al., 2024). Consider a face-to-face conversation: neural activity may align on linguistic features (e.g., the meaning of words) and non-linguistic features (e.g., gestures, facial expressions). To isolate *linguistic* features of shared brain activity across brains, we estimated encoding models from LLM embeddings (jointly with control features) and evaluated how well models trained on one subject generalize to their conversational partner. Specifically, given subject A and their conversational partner subject B, we correlated subject A’s production model predictions with subject B’s actual comprehension neural responses (Figure 5A). This analysis enabled us to test whether subject A’s encoding models in one conversational role can generalize and predict their partner’s neural responses in the other conversational role, vertex-by-vertex (Toneva et al., 2022; Zada et al., 2024). Our previous results showed that production and comprehension rely on similar brain regions and share similar linguistic features *within* subjects. This analysis reveals areas where production and comprehension are linguistically coupled *between* subjects.

**Figure 5.**
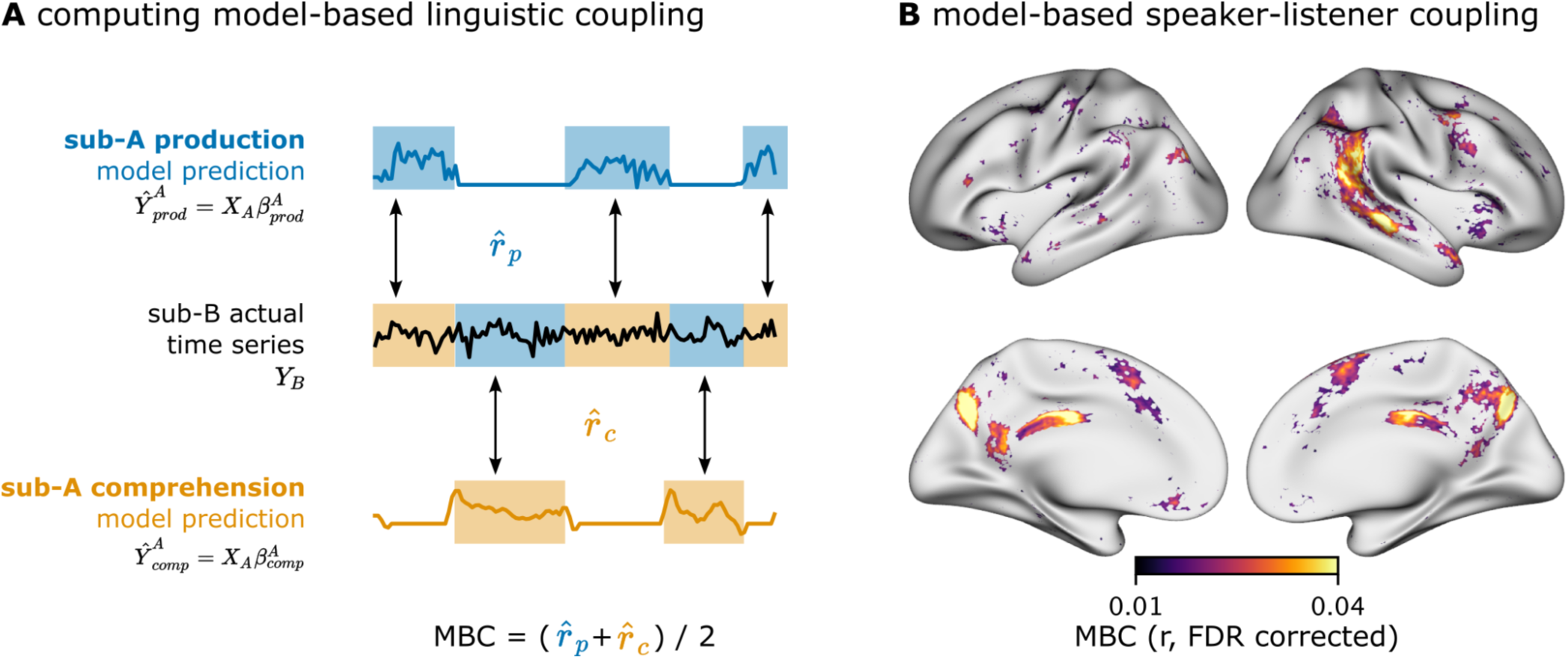
Model-based speaker-listener coupling. (**A**) Schematic of model-based coupling (MBC). We use the already-trained encoding models from subject A’s production data to predict subject B’s time series during comprehension. We correlate subject A’s model predictions with subject B’s time series separately for production and comprehension to obtain two correlations per subject per trial. (**B**) We average production and comprehension coupling correlations to obtain a group map of model-based coupling.

We found significant model-based speaker-listener coupling for LLM embeddings in the right hemisphere along pSTG, extending into the TPJ, the MFG, and bilaterally in precuneus in PMC (Figure 5B). Because the trained encoding model has to generalize to another subject’s brain performing a different process (production vs comprehension), the overall magnitude of the correlation is lower. Interestingly, this model-based linguistic coupling appears right-lateralized (in right-handed subjects). While relatively few vertices in left-hemisphere language areas were significant, we observed strong coupling in right-lateralized temporal areas and in bilateral PMC. For example, brain-to-brain coupling for LLM embeddings was found in the right TPJ, a structure commonly associated with mentalizing and social cognition (Frith & Frith, 1999, 2021). Thus, unlike within subjects where we find broad and bilateral model-based coupling (e.g., in STG, IFG, and PMC), model-based coupling between speaker and listener relies on right-lateralized pSTG and TPJ regions and bilateral precuneus, which are regarded as higher-order cognition areas.

### Spatial and temporal network structure in model-based conversational coupling

So far, we restricted the scope of speaker-listener coupling in both spatial and temporal dimensions for simplicity: we have only considered coupling between one brain area in the speaker and the homologous area in the listener, and we have only considered instantaneous, or “zero-lag” coupling between partners. The reality is much more complicated. For example, activity in some areas of the speaker’s brain may be coupled to activity in *different* regions of the listener’s brain, and in some cases, the speaker’s brain may *precede* that of the listener (Stephens et al., 2010; Zada et al., 2024). Here, we briefly explore variations in coupling along both of these axes. Since vertices are plentiful, adding spatial and temporal dimensions would exponentially increase the number of comparisons. Thus, we constrain this exploratory analysis to the 11 discussed ROIs.

We first assessed how well a model trained on the speaker’s brain activity in one ROI generalizes to the listener’s brain activity across all other ROIs. We did this by averaging the predicted and actual time series within each ROI across its vertices. This generated an inter-regional generalization matrix that summarizes the speaker-listener coupling across all ROI pairs at lag 0 (Figure 6A). We observed that the right hemisphere is more connected between speaker and listener than the left hemisphere. Moreover, some areas are relatively uncoupled from other regions (e.g., SM), whereas others are coupled with multiple areas (e.g., pSTG). Interestingly, this matrix has no clear diagonal, meaning that speaker-listener coupling across areas is similarly strong (or weak) to coupling between homologs.

**Figure 6.**
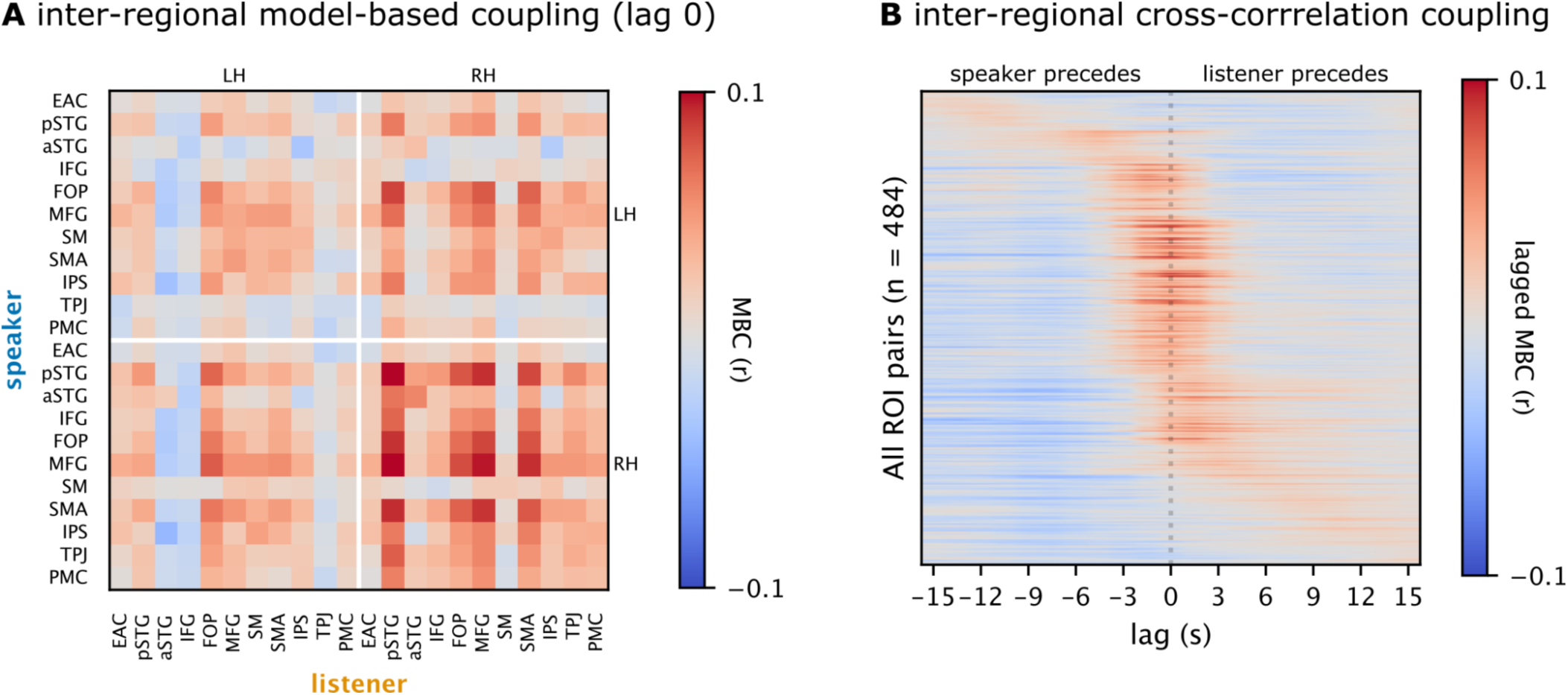
Inter-regional and cross-correlated model-based coupling. (**A**) We extend the speaker-listener model-based coupling results (Figure 5) along two dimensions. First, we correlate a speaker’s model-based prediction (averaged across vertices within ROIs) to all ROIs in the listener. (**B**) Second, for each pair of ROIs (a total of 484 pairs across both hemispheres), we cross-correlate the speaker’s predicted time series and the listener’s actual time series to extend coupling temporally. Rows are ordered by the lag at which they achieve maximum encoding performance.

To investigate temporal variation in linguistic coupling, we cross-correlated the predicted time series in subject A with the actual time series of subject B at varying lags (building off of Figure 5A). This resulted in a temporal profile for each ROI pair (484). A carpet plot of these profiles sorted by their peak lag suggests different clusters of temporal coupling (Figure 6B). Most pairs of regions exhibit peak coupling at lag 0 ± 3 seconds (e.g., TPJ, PMC, FOP). For a subset of region pairs, the speaker’s brain precedes the listener’s brain. For example, the speaker’s left MFG and SMA precede the listener’s brain activity (e.g., the speaker’s MFG precedes the listener’s pSTG). In another subset of region pairs, the listener’s brain appears to precede the speaker. For example, the speaker’s right aSTG and pSTG tend to lag behind the listener’s brain activity (e.g., the speaker’s aSTG lags behind the listener’s pSTG). These results suggest that linguistic coupling between conversation partners is spatially and temporally extended.

## Discussion

The neural systems involved in speech production and comprehension may require different processes, but they must converge on similar representations. After all, a shared linguistic space is necessary to align the linguistic information across the speaker’s and listener’s brains. This paper has sought to map the shared neural machinery mediating between speech comprehension and speech production in natural conversations. Our findings revealed that speech production and comprehension recruit similar brain areas and shared linguistic representations when engaged in natural conversation (Figure 2). In a test of generalization, we found that the brain’s linguistic processing during passive story listening is related to spontaneous speech production and comprehension during conversations (Figure 3). However, encoding performance was significantly weaker and missed key frontal language areas than when training on conversations. The model comparison analyses demonstrated that contextual embeddings from an LLM better capture the linguistic features shared between production and comprehension than other candidate models (Figure 4). Finally, our model-based coupling analysis revealed brain-to-brain production-comprehension coupling in high-level cortical areas, particularly right-hemisphere areas associated with social cognition (Figure 5). Overall, our results suggest that speech comprehension and speech production systems align on a set of shared, intermediate features, allowing the brain to translate between the two processes effectively.

We identified a unified language network with shared weights engaged during production and comprehension in real-time conversations. Encoding models have become essential for mapping linguistic features (e.g., acoustic, syntactic, and semantic features) to brain activity. Many recent studies have applied them during passive language comprehension (Caucheteux & King, 2022; de Heer et al., 2017; Deniz et al., 2019; Goldstein et al., 2022; Heilbron et al., 2022; Huth et al., 2016; Kumar et al., 2024; Schrimpf et al., 2021). However, only a handful of recent studies have begun leveraging encoding models for spontaneous language production and active comprehension (Cai et al., 2025; Goldstein et al., 2025; Yamashita et al., 2025), and even fewer have simultaneously recorded two participants engaged in dialogue (Spiegelhalder et al., 2014; Zada et al., 2024). Our encoding models were able to predict neural responses during spontaneous speech production and comprehension (Figure 2A). They also provide an elegant way of comparing these processes within subjects. By constraining the model to share weights, we found that most brain regions exhibited shared linguistic representations between production and comprehension (Figure 2B). Thus, providing evidence for an overlap between speech production and comprehension, which relies on a unified and shared language network. Part of this common network constitutes well-established language regions (Fedorenko et al., 2024), and extends into general systems responsible for interactive, social cognition. We also found a production-comprehension overlap in low-level perceptual and motor areas (e.g., EAC, SM), suggesting that modality-specific areas may be more localized than previously thought. By using natural conversations, we were able to demonstrate how participants engage these neural processes in real-world, interactive communication (Hagoort, 2019; Wheatley et al., 2024) that embody the principles of ecological validity in social neuroscience (Hasson & Honey, 2012; Nastase et al., 2020; Zaki & Ochsner, 2009).

Advances in simultaneous neuroimaging allowed us to move beyond asynchronous protocols of speaker-listener coupling to real-time, turn-taking conversations. Our hyperscanning paradigm allowed us to simultaneously record brain activity during speech production and speech comprehension in two interacting subjects. Whereas landmark studies were limited to relating a single subject’s production to multiple subject’s comprehension responses acquired at a later time (Silbert et al., 2014; Stephens et al., 2010), our paradigm engages each subject’s production and comprehension processes in an (inter)active, real-time, turn-taking conversation. We found that conversations recruit more brain regions and different representations than non-interactive paradigms (Figures S1, S5). Similar to studies of asynchronous communication, we observed production-comprehension coupling in PMC, pSTG, and TPJ (Figure 5B). While these studies relied on a content-agnostic approach to speaker-listener coupling (e.g., using ISC, see Introduction), we used a model-based approach to quantify *linguistic* coupling across speakers. Rather than merely showing us *where* coupling occurs, this approach allows us to explicitly model *what* features are coupled across brains (Zada et al., 2024). Moreover, by explicitly representing different linguistic features in one model, we ensure that it is the linguistic information from the contextual LLM embeddings that we find coupled within and between the subject’s production and comprehension processes (Figure 4). In doing so, this approach does not register coupling in EAC, as Stephens and colleagues (2010) reported, which likely stems from synchronous, low-level auditory speech features, rather than contentful representations.

We speculate that interactive communication, where partners must actively listen and figuratively “speak to” one another’s thoughts and intentions, may engage the social brain in a way that traditional language paradigms do not. Historically, language processing—both comprehension and production processes—has been associated with the left hemisphere (Broca, 1865; Corballis, 2014; Dax, 1865; Knecht et al., 2000; Wernicke, 1874). On the other hand, both ISC analyses (e.g., Nastase et al., 2021) and encoding models (e.g., Huth et al., 2016) tend to yield largely bilateral maps during natural language comprehension. In the current study, we observed brain-to-brain linguistic coupling in the right-lateralized superior temporal cortex, TPJ, and prefrontal cortex, as well as bilateral precuneus and posterior cingulate. This result indicates that the same features that mediate between comprehension and production processes within a brain are also partly shared across individuals. However, these areas are not simply right-hemisphere homologs of typical language regions (Braga et al., 2020; Fedorenko et al., 2024). In the neuropsychology literature, the right hemisphere has been associated with affective and other paralinguistic features of speech (Heilman et al., 1975; Lindell, 2006), as well as pragmatic and discourse-level processing (Beeman, 1993; Beeman & Chiarello, 1998; Kaplan et al., 1990). Neuroimaging work has generally corroborated these findings (Bottini et al., 1994; Gernsbacher & Kaschak, 2003; Robertson et al., 2000; Vigneau et al., 2011); for example, Yarkoni and colleagues (2008) reported a very similar set of regions to ours, including right TPJ, and bilateral posterior cingulate and precuneus, involved explicitly in tracking narrative comprehension across sentences. Interestingly, several of these areas overlap with regions often associated with mentalizing and other aspects of social cognition (Frith & Frith, 2012; Saxe, 2006), highlighting the key role that the social brain may play in real-time, naturalistic social interactions.

Large language models are trained to predict the next word in large text corpora. After training, these models can generate increasingly fluent, surprisingly meaningful language, one word at a time, by sampling from a probability distribution of upcoming words. These models do not have dedicated systems for comprehension or production resembling anything like the human brain. Why do these models capture neural activity so well during language comprehension and production? Generative language models operate in a simple perception-action loop by mapping each current word to predict the upcoming word (Pulvermüller, 2018). We speculate that this constraint, which forces language models to learn *shared* representations that inform upcoming word predictions, may yield embeddings that can capture brain activity during both comprehension and production. While the brain has specialized systems for perception and production, our findings suggest that many of the brain’s language machinery occupies a middle ground to LLM embeddings: multimodal, active representations with mixed features for both comprehension and production.

## Methods

### Participants

Thirty dyads (*N* = 60 participants) engaged in real-time conversations while they were simultaneously scanned with fMRI hyperscanning. These data are a subset of a larger dataset collected with additional conditions and participants (see Speer et al., 2024). Participants were recruited from Princeton University and received monetary compensation for their participation. Eligibility requirements included: must be 18 years or older, right-handed, and with normal or corrected vision. Of the 58 included participants, 41 were female, and the average age was 20.74 (minimum 18, maximum 36). One dyad was excluded due to an unexpected scanning issue that resulted in fewer conversations than others.

### Design

Two participants at a time arrived at two fMRI scanners in adjacent rooms. The participants did not know each other before the experiment but briefly met before entering the scanners. Participants were instructed to engage in prompted conversations across five runs. Prompts were specifically designed to increase the level of intimacy of conversations across the runs, and are based on stimuli from Aron and colleagues (1997) (Table S1). Each run was 13:36 minutes long and consisted of four trials. We only used two trials of each run because the other two trials were not spontaneous conversations, and were used for a different experiment. Each trial was 03:21 minutes long and started with the topic prompt displayed on screen for 9 seconds, followed by the conversation for 180 seconds, and ended with 12 seconds of a fixation cross (Figure 1). The participant who would start speaking first was randomly assigned. Once a participant finished their utterance, they were instructed to press a button to “pass the virtual mic” to their conversational partner. When a participant had the virtual mic, the screen displayed the text “your turn to speak, when you want to pass the mic, press ‘1’”, followed by a countdown timer displaying the number of seconds left. When listening, the screen showed “your turn to listen”, followed by the same countdown timer. Participants were instructed to fill the entire three minutes. After all runs, participants filled out a survey answering questions about the level of enjoyment, similarity, and closeness they felt during their conversations.

### MRI acquisition

We recorded neuroimaging data using 3T Siemens Skyra and 3T Siemens Prisma MRI systems. Both machines were configured using the same scanning parameters. Functional scans were acquired with whole brain coverage in interleaved order: 3.0 mm slice thickness, 3.0 × 3.0 mm in-plane resolution, flip angle = 80°, TE = 28 ms, TR = 1500 ms, multiband acceleration factor = 2. A T1-weighted image was acquired for anatomical reference: 1.0 × .0 × 1.0 mm resolution, 176 sagittal slices, flip angle = 9°, TE = 2.98 ms, TR = 2300 ms. To minimize head movement, the subjects’ heads were stabilized with foam padding.

### Conversation audio transcription

Each three-minute audio segment was transcribed, aligned, and diarized (assigned unique speaker labels) at the word level using WhisperX (Bain et al., 2023)—an automatic speech recognition tool. We used the *faster-whisper-large-v2* model and set the minimum and maximum speakers to two. Each resulting transcription consisted of each word spoken, its onset and duration, and the identity of the speaker.

### fMRIPrep preprocessing

Results included in this manuscript come from preprocessing performed using *fMRIPrep* 20.2.0 (Esteban et al., 2018, 2019), which is based on *Nipype* 1.5.1 (K. Gorgolewski et al., 2011; K. J. Gorgolewski et al., 2018) and *Nilearn* 0.6.2 (Abraham et al., 2014).

T1-weighted images were corrected for intensity non-uniformity (INU) with *N4BiasFieldCorrection* (Tustison et al., 2010), distributed with ANTs 2.3.3 (Avants et al., 2008), and used as a reference throughout the workflow. The T1 reference was then skull-stripped with a *Nipype* implementation of the *antsBrainExtraction.sh* workflow (from ANTs), using OASIS30ANTs as target template. Brain tissue segmentation of cerebrospinal fluid (CSF), white-matter (WM) and gray-matter (GM) was performed on the brain-extracted T1 image using *fast* (FSL 5.0.9 Zhang et al., 2001). Brain surfaces were reconstructed using *recon-all (FreeSurfer 6.0.1 Dale et al., 1999)*, and the brain mask estimated previously was refined with a custom variation of the method to reconcile ANTs-derived and FreeSurfer-derived segmentations of the cortical gray-matter of Mindboggle (Klein et al., 2017). Individual cortical surface reconstructions were aligned to the *fsaverage6* surface template (40,962 vertices per hemisphere) based on sulcal curvature (Fischl et al., 1999).

### Functional data preprocessing

For each of the 6 BOLD runs found per subject (across all tasks and sessions), the following preprocessing was performed. First, a reference volume and its skull-stripped version were generated. A deformation field to correct for susceptibility distortions was estimated based on *fMRIPrep*’s *fieldmap-less* approach. The deformation field is constructed by co-registering the BOLD reference to the same-subject T1 reference with inverted intensity (Huntenburg, 2014; Wang et al., 2017). Registration is performed with *antsRegistration* (ANTs 2.3.3), and the process is regularized by constraining deformation to be nonzero only along the phase-encoding direction, and modulated with an average fieldmap template (Treiber et al., 2016). Based on the estimated susceptibility distortion, a corrected BOLD reference was calculated for a more accurate co-registration with the anatomical reference.

The BOLD reference was then co-registered to the T1w reference using FreeSurfer’s *bbregister*, which implements boundary-based registration (Greve & Fischl, 2009). Co-registration was configured with six degrees of freedom. Head-motion parameters with respect to the BOLD reference (transformation matrices, and six corresponding rotation and translation parameters) were estimated before any spatiotemporal filtering using *mcflirt* (FSL 5.0.9 Jenkinson et al., 2002). BOLD runs were slice-time corrected using *3dTshift* from AFNI 20160207 (Cox & Hyde, 1997). The BOLD time series were ultimately resampled onto the *fsaverage6* surface template using FreeSurfer’s *mri_vol2surf*. Resampling was performed with a single interpolation step by applying a single, composite transform to correct for head motion, slice-timing, susceptibility distortions, and normalization to the surface template. All subsequent analyses were applied to the vertex-level functional data in surface space; our use of the term “vertex” is otherwise synonymous with the use of “voxel” in volumetric analyses (e.g., “voxelwise encoding models”).

Several confounding time series were calculated while preprocessing the BOLD data: six head motion parameters, framewise displacement (FD), and a set of physiological components. FD was estimated for each functional run by computing the absolute sum of relative motions (Power et al., 2014). FD was calculated for each functional run using the implementation in *Nipype* (following the definitions by Power et al., 2014). The three global signals are extracted within the CSF, the white matter, and the whole-brain masks. Additionally, a set of physiological regressors were extracted to allow for anatomically constrained component-based noise correction (aCompCor Behzadi et al., 2007). Principal components are estimated after high-pass filtering the preprocessed BOLD time series using a discrete cosine filter with 128s cut-off. We retained 10 aCompCor components, five estimated from a white matter mask, and five from a CSF mask.

### Confound regression and head motion correction

A typical fMRI signal cleaning pipeline involves regressing out nuisance variables from fMRIPrep’s output from the BOLD signal across an entire run or scan (Ciric et al., 2017; e.g., Friston et al., 1996; Parkes et al., 2018; Satterthwaite et al., 2013). Nuisance variables include head motion (e.g., rigid-body motion parameters), physiological noise (e.g., cardiac fluctuations), and scanner noise (e.g., signal drift). However, our hyperscanning paradigm with freely alternating speech production and comprehension between subjects requires additional task-related nuisance variables.

From fMRIPrep confounds, we chose the six head motion variables, all available cosine variables, and the top five components from aCompCor for white matter and CSF masks, separately. This resulted in 26 nuisance regressors. Next, we added five regressors based on the task structure (see the previous Design section). Three boxcar regressors were initialized with zeros across the entire run and populated with ones for (1) indicating the two different trial types, (2) indicating turn to speak, and (3) indicating turn to listen. Two indicator regressors were initialized with zeros and filled with ones when either (1) the subject pressed the button to end their turn, or (2) their conversation partner pressed the button (the instructions on the screen switched each time a button was pressed). These regressors were convolved with an HRF to account for the hemodynamic response using Nilearn’s *glm.first_level.glover_hrf* implementation. Finally, all confound variables were passed to Nilearn’s *signal.clean* function to detrend, regress out the variables, and z-score the time series.

### Defining cortical regions of interest

In order to summarize results across the cortex, we first aggregated the 40,962 vertices in each hemisphere into 180 parcels from a widely-used Glasser multimodal parcellation (Glasser et al., 2016). Then, we defined an extended parcel-level language network from four primary sources: a collection of functionally defined language regions (Fedorenko et al., 2010), a probabilistic atlas based on language localizer tasks in 806 subjects (Lipkin et al., 2022), an activation map corresponding to the “language” topic from NeuroSynth (Yarkoni et al., 2011), and an intersubject correlation map (ISC) based on 345 subjects listening to natural stories (Nastase et al., 2021). We thresholded the probabilistic atlas at *p*=0.10, the NeuroSynth map at *t*=0.10, and the intersubject map at *r*=0.10. We overlaid these four maps to form an extended “meta” map of language areas (Figure S3A).

We grouped the 55 parcels within this final brain map into 11 regions of interest based on their spatial proximity and previously identified groupings (Figure S3B, Table S2). Specifically, following the networks identified by Glasser and colleagues (2016), we identified the following regions: early auditory cortex (EAC), posterior and anterior superior temporal gyrus (pSTG, aSTG), inferior and middle frontal gyri (IFG, MFG), somatomotor cortex (SM), supplementary motor area (SMA), frontal operculum (FOP), intraparietal sulcus (IPS), temporoparietal junction (TPJ), and posterior medial cortex (PMC). Finally, given that the maps derived from prior studies may be biased toward comprehension tasks, we defined a somatomotor region of interest we expect to be involved in language production (Silbert et al., 2014). Note that we are deliberately defining a more inclusive “language network” than prior work (Fedorenko et al., 2010) to explore both more peripheral perception (e.g., EAC) and production (e.g., SM) areas, as well as higher-level areas that may be involved in narrative and social cognition (e.g., TPJ, PMC).

### Linguistic features for encoding analysis

In vertex-wise encoding analysis, we use ridge regression to learn a linear model mapping from a set of explicit features (i.e., design matrix) to the observed brain activity (Naselaris et al., 2011). We first re-represent the language task and stimulus in one or more feature spaces. We defined several feature spaces from the conversation stimuli to build these design matrices.

#### Task structure and nuisance variables

We computed four low-level variables from each transcript that could affect the BOLD signal (Huth et al., 2016). For each TR, we quantify the word rate (number of words in a TR), phoneme rate (number of phonemes in a TR), word occurrence (some TRs contained no words), and a variable indicating whether it was the subject’s turn to speak or listen. The word and phoneme rates were continuous, while the word onset and indicator variables were binary.

#### Acoustic spectral features

For each pair of subjects, we had one audio recording of the entire conversation that was recorded from one mic at a time and switched upon button presses indicating the end of turn. We computed acoustic features from the speech audio files (de Heer et al., 2017). Specifically, we used the *WhisperFeatureExtractor* class from the *HuggingFace* (Wolf et al., 2020) library with the default settings to extract a spectral representation of the audio. This function uses a short-time Fourier transform to compute a mel-filter bank of 80 features that represent the spectral power density on a Mel log scale. Note that these features likely capture more than just acoustic features because they were recorded in MRI machines with different noise characteristics, and were saved into one file from two sources. Thus, at minimum, it also encodes information about the conversation turns.

#### Articulatory phonemic features

Following de Heer et al., (2017), we quantify the articulatory features of speech based on the phonemes in the transcript. Specifically, we used the CMU pronunciation dictionary (http://www.speech.cs.cmu.edu/cgi-bin/cmudict) to obtain the phonemes associated with each word in the transcript. We then constructed the articulatory features for each phoneme based on the place and manner of consonants, and voicing of vowels. This resulted in a binary vector of 22 features for each phoneme.

#### Large language model features

We extracted word embeddings from the large language model GPT-2 XL (Radford et al., 2019) using the *HuggingFace* library (Wolf et al., 2020). For each 3-minute conversation transcript, we first converted all words to GPT-2 tokens. We then passed these tokens as input to the LLM, where they were converted to 1,600-dimensional token embeddings and passed through the decoder layers. We extracted the activations from the middle (24th) layer to serve as contextual word embeddings.

### Encoding model construction and evaluation

Encoding models were the core analytical approach we took to estimating linguistic content in the BOLD signal (Naselaris et al., 2011). For all analyses, we used kernel ridge regression to prevent overfitting, and banded ridge regression to find different regularization parameters for each feature space separately (Nunez-Elizalde et al., 2019). We used the *MultipleKernelRidgeCV* class from the *himalaya* library (Dupré La Tour et al., 2022) to perform cross-validation within the training set to select the best regularization parameter per feature space. All results we report on encoding performance were evaluated on a held-out test sample.

#### Design matrix construction

Each 3-minute conversation (trial) consisted of a 120-TR BOLD time series. With two trials per run (240 TRs) and five total runs, we had a total of 1,200 TRs per subject. Thus, our design matrix had 1,200 rows. The initial number of columns was based on the selected feature spaces for each analysis. For example, for the full joint model (Figure 1), we used five feature spaces: task (8 dimensions), acoustic (80), articulatory (22), and contextual embeddings (1,600). Stimuli features that were defined on the word or token level were averaged within TRs (e.g., LLM embeddings). Then, we split each feature space into two groups, for production or comprehension, and filled the gaps between one process and the other with zeros.

#### Model definition

We used a Scikit-learn (Pedregosa et al., 2011) pipeline to define the full encoding model. The pipeline consisted of three main steps before model fitting. First, the regressors were mean-centered using *StandardScaler*. Then, each feature space was duplicated and shifted by 2–5 TRs (3–7.5 s) to account for the hemodynamic lag in the BOLD signal (Huth et al., 2016). Finally, because the design matrix was wider than it is long, we used the kernel method to solve the ridge regression in its dual form (Dupré La Tour et al., 2024). Specifically, we used a linear kernel for each feature space separately before fitting the model.

#### Model fitting and evaluation

We used cross-validation to evaluate each model on a held-out test sample. Specifically, we defined five folds, based on the five runs, to fit a model on four runs (960 TRs), and tested it on the held-out run (240 TRs). We repeated this procedure five times, testing each run in turn, and then averaging the encoding performance across the five runs. Each run contained unique conversations based on different prompts.

Banded ridge regression allows us to evaluate each feature space separately, relative to all the others. To do this, the joint predicted time series on the held-out run can be decomposed into one time series per feature space (Dupré La Tour et al., 2022). Similarly, the encoding model performance (i.e., the correlation between the predicted and actual time series) can be split into one correlation for each feature space. Importantly, we segmented the actual and predicted time series into production and comprehension TRs to obtain their separate correlations for each process. Moreover, because of the hemodynamic response, some TRs may be affected by both processes. Thus, we selected the exclusive set of TRs where there is no overlap.

Finally, we confirmed that head motion degrades encoding performance and that there is considerably more head motion during speech production than comprehension (Figure S4).

#### Statistical significance

We tested whether a vertex’s encoding performance correlation is statistically significant by using a two-sided, one-sample *t*-test, as implemented in SciPy (Virtanen et al., 2020). All p-values were corrected for multiple comparisons by controlling the false discovery rate (FDR; Benjamini & Hochberg, 1995).

### Speaker–listener model-based coupling

We used the already-trained encoding models to evaluate the model-based coupling between conversation partners. The intuition behind this evaluation is to correlate one subject’s model-predicted time series with their conversational partner’s actual time series (as opposed to correlating it with their *own* actual time series). In effect, this simultaneously tests whether the model can generalize from one subject to another and from one process to another (e.g., production to comprehension) (Toneva et al., 2022). Thus, we use the same evaluation procedure as described before, except with one major change. For each voxel, we correlate a subject’s predicted time series with their partner’s actual time series for the same voxel. Critically, we use the predictions from all feature spaces and compute the relative encoding performance of the LLM contextual embedding feature space only. By applying the same evaluation procedure as within-subject, we control for variance that can be explained by the nuisance feature spaces. When testing model-based coupling across regions and time (Figure 6), we first extract speaker turns that are at least 9 seconds long in order to exclude turns that are too short.

### Story-listening task and analysis

Prior to hyperscanning acquisition, participants listened to a ∼13-minute story (“I Knew You Were Black” by Carol Daniel). Three participants did not complete this task and were excluded from this particular analysis. We used the same procedures as described above for conversations for the story, including MRI acquisition parameters, BOLD preprocessing, confound regression, linguistic features, and encoding model construction, training, and evaluation. However, there were two differences. First, the confound regression did not include any design structure variables. Second, we did not split regressors because the story is only comprehension—thus this model corresponds to the shared-weights model for conversations.

### Software resources

In addition to the software mentioned throughout the Methods, we used *Surfplot* (Gale et al., 2021) for visualizing brain maps.

## Acknowledgments

We would like to extend thanks to Ahmad Samara and Itamar Jalon for helpful feedback on head motion correction and fMRI preprocessing details.

## Author contributions

Conceptualization: ZZ, SAN, UH, DT

Data curation: LT, SB, SS

Formal analysis: ZZ

Funding acquisition: DT

Investigation: ZZ

Methodology: ZZ

Project administration: DT

Software: ZZ

Supervision: SAN, DT

Visualization: ZZ

Writing – original draft: ZZ

Writing – review & editing: ZZ, SAN, LMT, UH, DT

## Data and materials availability

Code for all results in this manuscript is publicly available on GitHub (https://github.com/zaidzada/fconv). Neural data and transcripts not currently available to protect participant privacy.

