## Supplementary Materials for "Linguistic coupling between neural systems for speech production and comprehension during real-time dyadic conversations"

Includes supplementary figures S1–S4 and tables S1–S2.

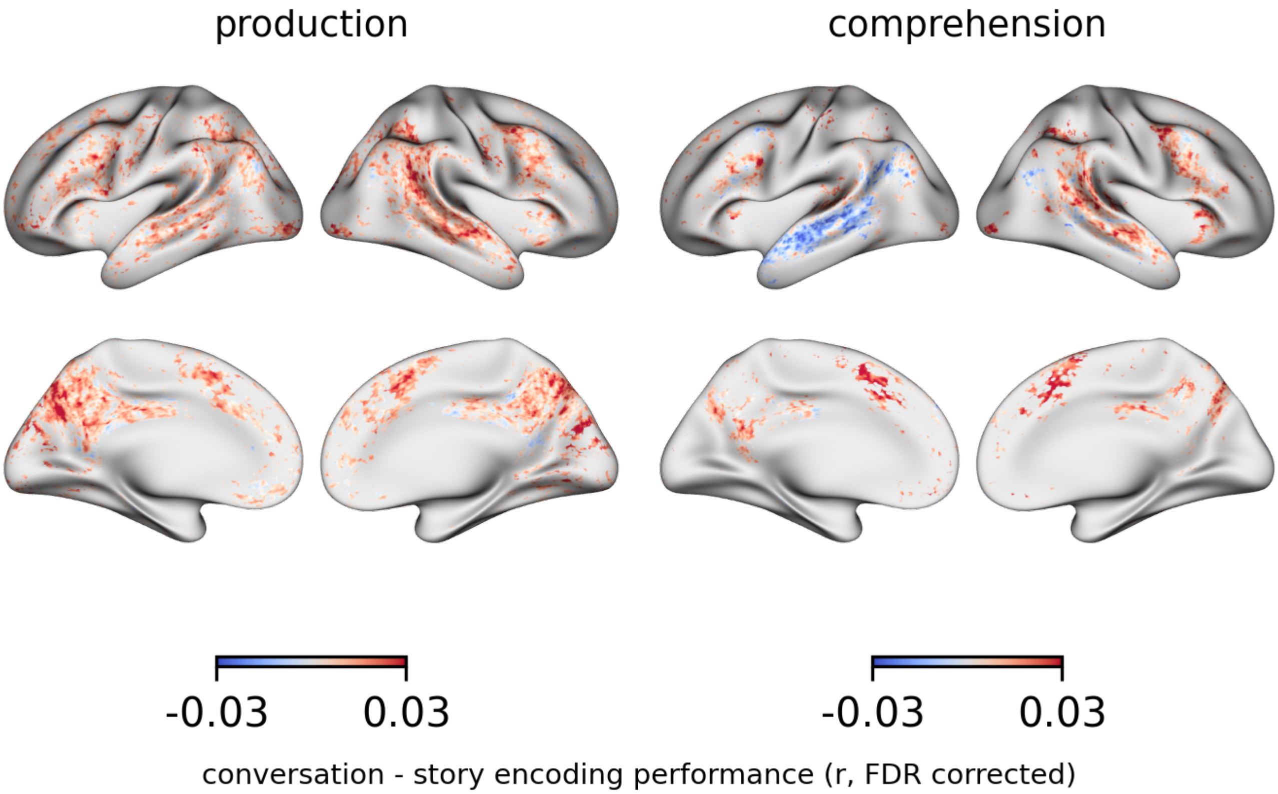


**Figure S1. Within-subject contrast between training on conversations versus story.** To fairly compare encoding performance between training encoding models on conversational data or story listening data, we performed an additional analysis. Instead of the 5-fold cross validation procedure used before, here we held out 3 conversation runs for testing for both story- and conversation-trained models. For the story, we trained encoding models on all 534 TRs of the story, while for the conversation, we trained encoding models on the first two runs only (480 TRs of both production and comprehension). For conversational encoding models, we used the shared weights model. Thus, this analysis ensures that both encoding models are tested on the same data and have roughly similar training set sizes. Evaluation of each model was performed *within-subjects* (i.e., training and testing on each subject's data separately). We plot the difference between conversation- and story- encoding model performance while thresholding for significantly predicted vertices only. Positive numbers (red) reflect vertices where training on conversational data performs better, and vice versa for negative numbers (blue).

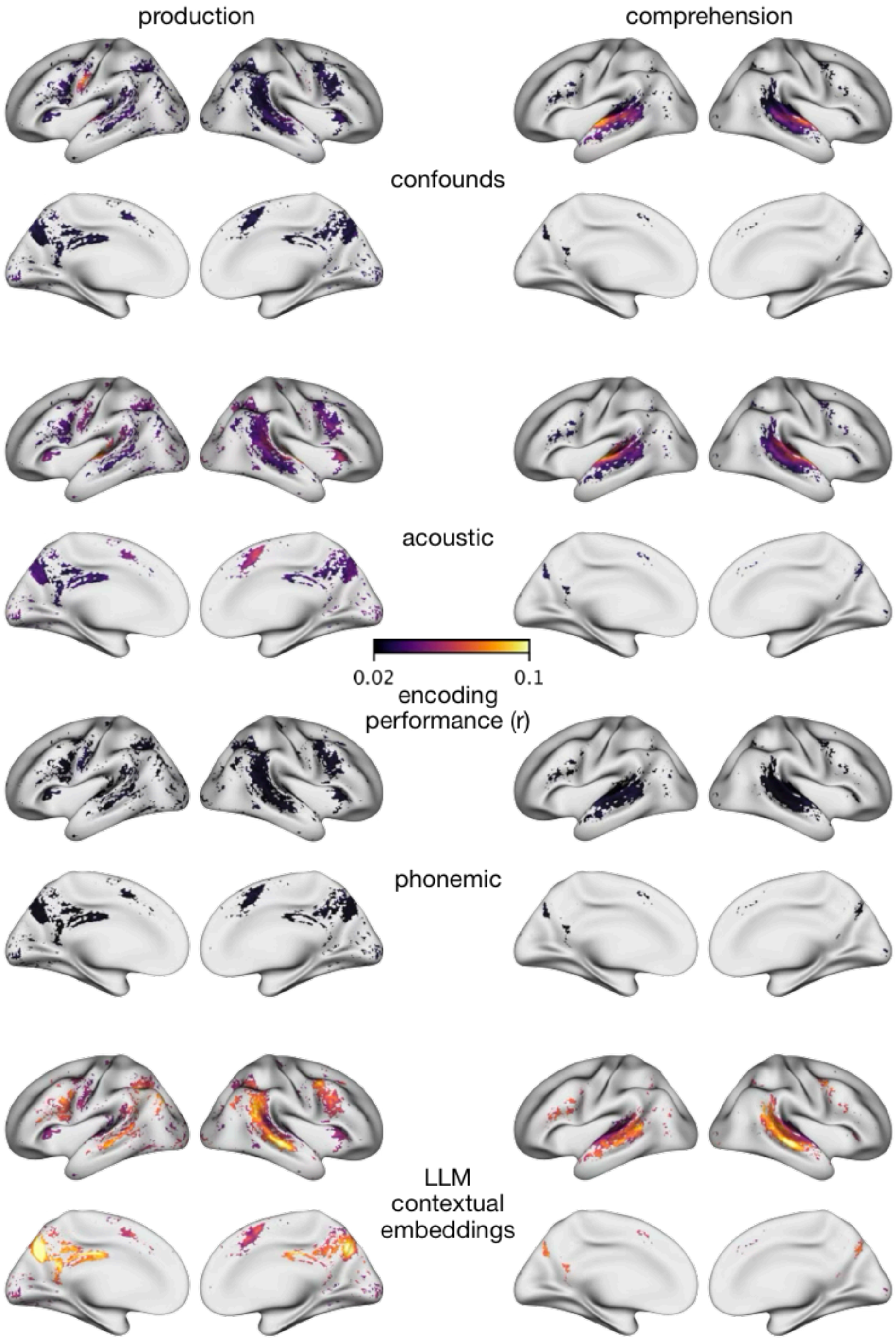


**Figure S2. Within-subject encoding performance per band during production and comprehension.** The joint model encoding performance can be decomposed into the relative contribution per feature space (Figure 1). Moreover, we evaluate production time points separately from comprehension time points. Here, we threshold the brain maps using a one-sample *t*-test based on the joint model performance, and then apply Bonferroni correction.

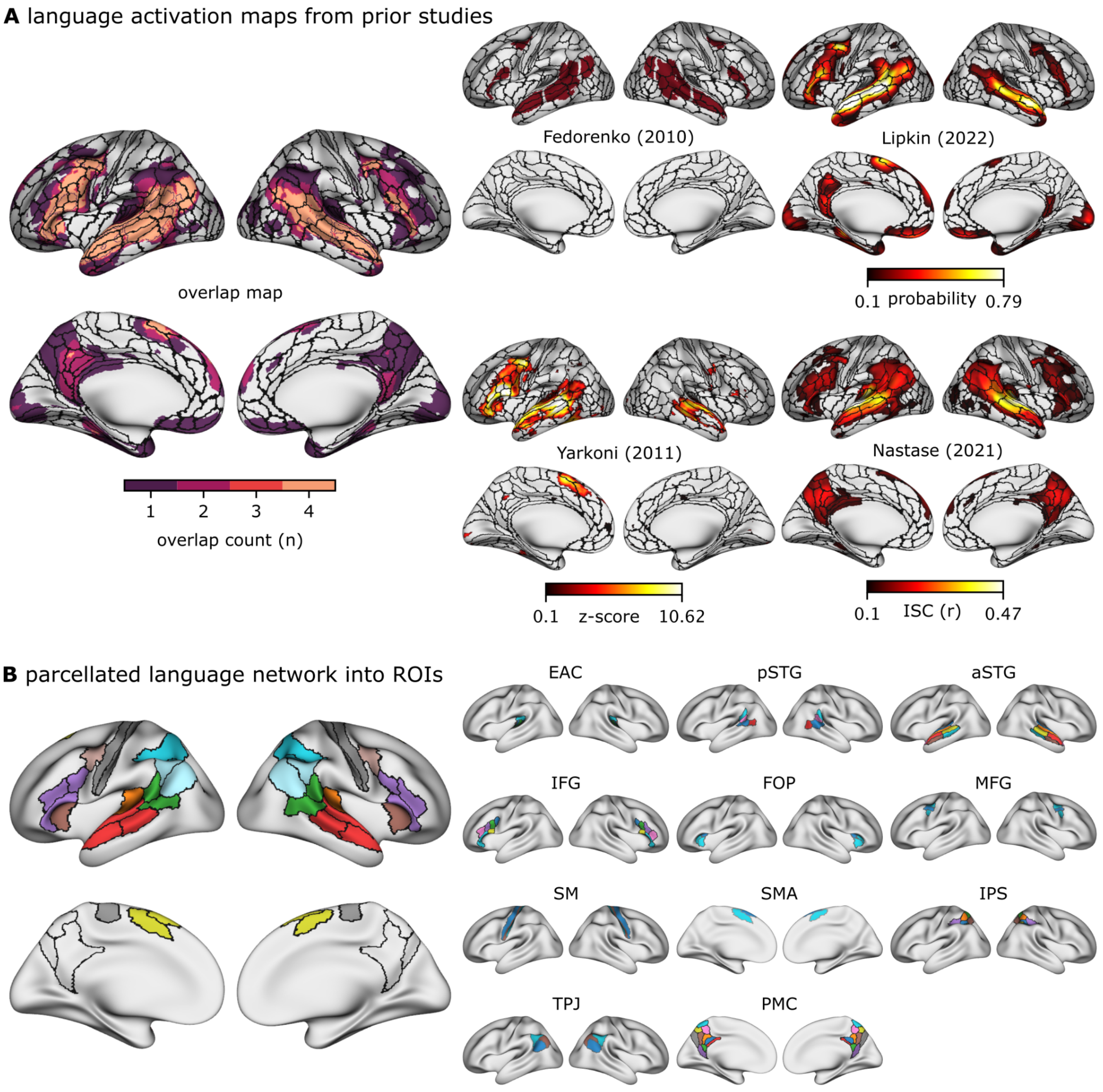


**Figure S3. Regions of interest within the extended language network.** (**A**) We define linguistic regions of interest based on the overlap of four primary sources of language-related brain maps. See Methods for details on thresholding. (**B**) Then, we select parcels in the Glasser atlas where overlap occurs, and group parcels into 11 regions per hemisphere.

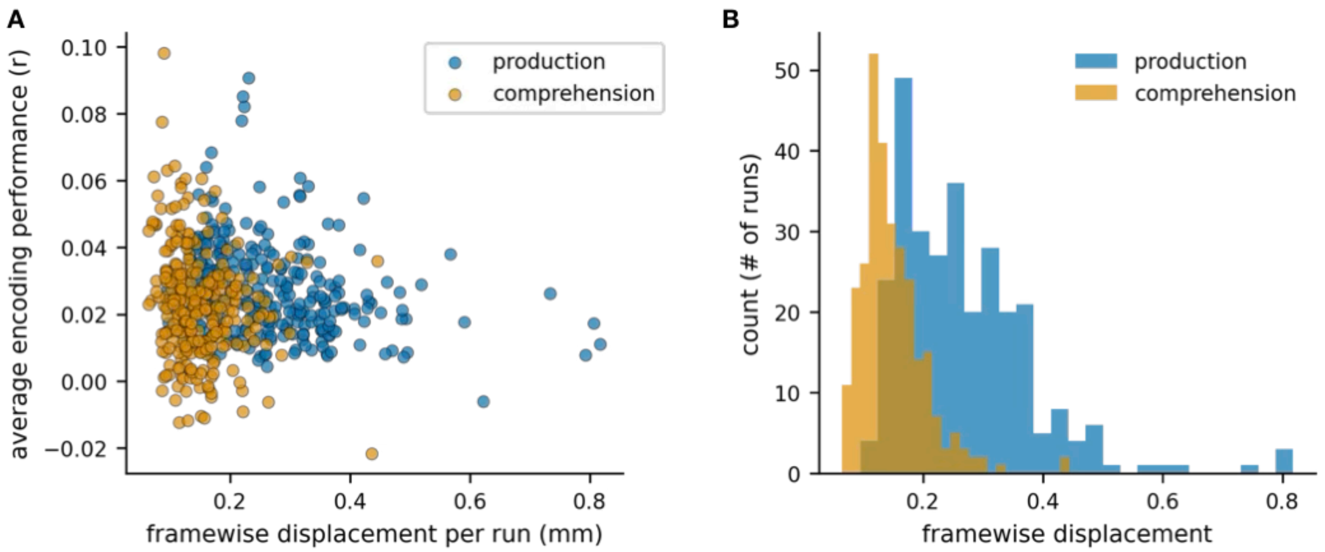


**Figure S4. Head motion impact on model performance.** (**A**) We found that head motion degrades the model performance for both production ($r = -0.291$, $p < 1e\text{-}07$) and comprehension ($r = -0.207$, $p < 0.00038$). (**B**) Production and comprehension histogram of the average framewise displacement per subject for each of their five runs. As expected, more head motion (as measured with framewise displacement) occurs during production than comprehension.

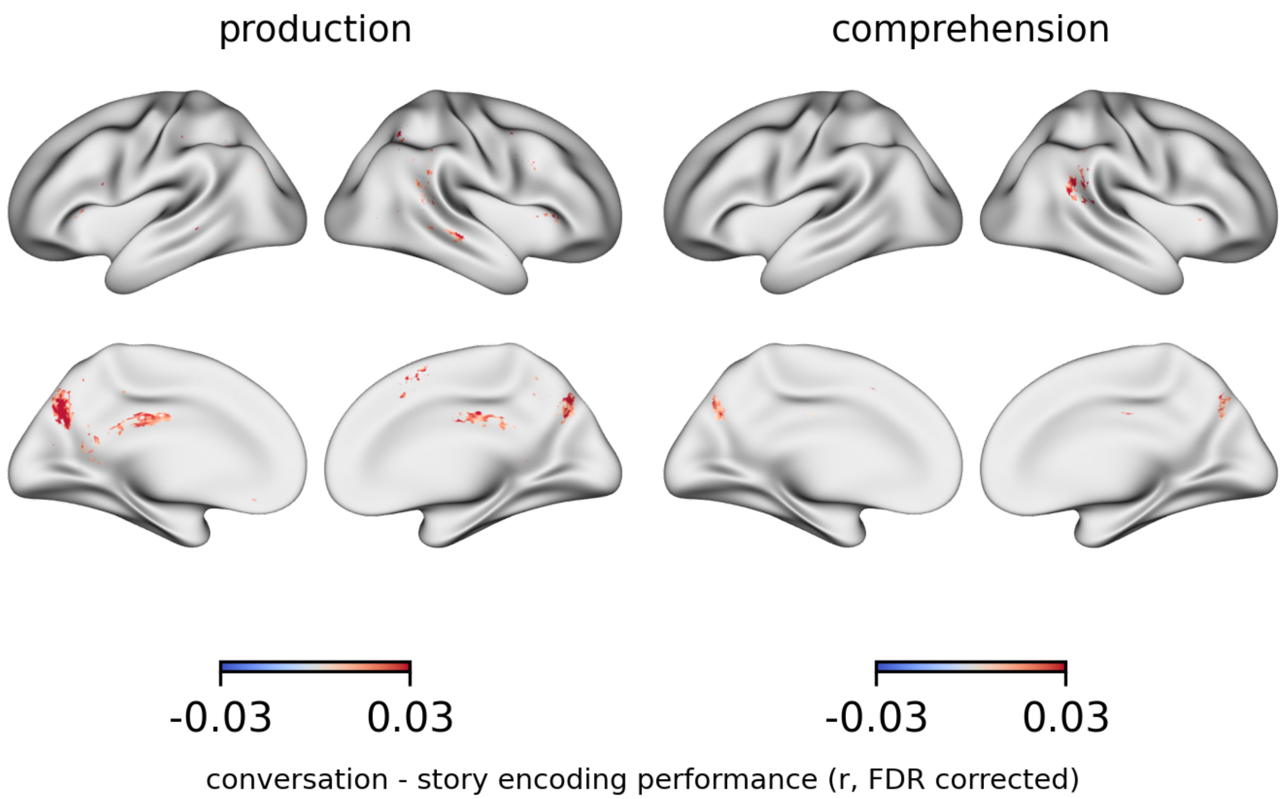


**Figure S5. Across-subject contrast between training on conversations versus story.** Here we test the difference in encoding performance *across subjects* when training on conversational or story data. The procedure is the same as Figure S1 but instead of evaluating each model on a subject's own data, we test it on their conversational partner's neural data for the three held-out conversation runs.

| prompt | set | prompt text |
|---|---|---|
| 1 | 1 | Given the choice of anyone in the world, whom would you want as a dinner guest? |
| 2 | 1 | Would you like to be famous? In what way? |
| 3 | 1 | Before making a telephone call, do you ever rehearse what you are going to say? Why? |
| 4 | 1 | What would constitute a "perfect" day for you? |
| 5 | 1 | When did you last sing to yourself? To someone else? |
| 6 | 1 | If you were able to live to the age of 90 and retain either the mind or body of a 30-year-old for the last 60 years of your life, which would you want? |
| 7 | 1 | For what in your life do you feel most grateful? |
| 8 | 1 | If you could change anything about the way you were raised, what would it be? |
| 9 | 2 | If a crystal ball could tell you the truth about yourself, your life, the future, or anything else, what would you want to know? |
| 10 | 2 | Is there something that you've dreamed of doing for a long time? Why haven't you done it? |
| 11 | 2 | What is the greatest accomplishment of your life? |
| 12 | 2 | What do you value most in a friendship? |
| 13 | 2 | What is your most treasured memory? |
| 14 | 2 | How close and warm is your family? Do you feel your childhood was happier than most other people's? |
| 15 | 3 | Complete this sentence: I wish I had someone with whom I could share… |
| 16 | 3 | Please share what would be important for your study partner to know as your close friend. |
| 17 | 3 | Share with your partner an embarrassing moment in your life. |
| 18 | 3 | What, if anything, is too serious to be joked about? |
| 19 | 3 | Your house, containing everything you own, catches fire. After saving your loved ones and pets, you have time to safely make a final dash to save any one item. What would it be? Why? |
| 20 | 3 | Share a personal problem and ask your partner's advice on how he or she might handle it. Also, ask your partner to reflect back to you how you seem to be feeling about the problem you have chosen. |

**Table S1. Conversation topic prompts.** Participants were presented with 20 different prompts to inspire otherwise free-form conversations. The prompts were constructed to become increasingly personal over the course of the experiment.

| ROI | sub-group | parcels from Glasser (2016) |
|---|---|---|
| EAC | EAC | [A1, LBelt, MBelt, PBelt, RI] |
| pSTG | AG | [TPOJ1, TPOJ2] |
| pSTG | SMG | [STV, PSL] |
| aSTG | STG | [A4, A5] |
| aSTG | aSTS | [STSda, STSva, STGa] |
| aSTG | pSTS | [STSdp, STSvp] |
| IFG | IFJ | [IFJp, IFJa] |
| IFG | IFS | [IFSp, IFSa] |
| IFG | IFG | [44, 45, 47l] |
| MFG | MFG | [55b, FEF, PEF] |
| SM | M1 | [4] |
| SM | S1 | [3a, 3b] |
| FOP | FOP | [FOP4, FOP5, AVI] |
| SMA | SFL1 | [SFL] |
| SMA | SFL2 | [SCEF] |
| IPS | SPC | [LIPd, LIPv, VIP, AIP, MIP] |
| IPS | dIPC | [IP0, IP1, IP2] |
| TPJ | IPC1 | [PGi, PGs] |
| TPJ | IPC2 | [PFm] |
| PMC | PCC1 | [31pv, 31pd, v23ab, d23ab, POS1, 7m, PCV] |
| PMC | PCC3 | [POS2] |
| PMC | SPC1 | [7Pm, 7Am] |

**Table S2. Language network atlas constituents.** We constructed 11 ROIs spanning an extended language network, including early auditory areas, language areas, higher-level areas associated with semantic representation and narrative processing, and somatomotor areas. The ROIs were selected based on prior work (Figure S3) and constructed by combining parcels from a multimodal atlas (Glasser et al., 2016).
